# Ability of Bicarbonate Supplementation to Sensitize Selected Methicillin-Resistant *Staphylococcus aureus* (MRSA) Strains to β-Lactam Antibiotics in an *Ex Vivo* Simulated Endocardial Vegetation Model

**DOI:** 10.1101/807255

**Authors:** Warren E. Rose, Ana M. Bienvenida, Yan Q. Xiong, Henry F. Chambers, Arnold S. Bayer, Selvi C. Ersoy

## Abstract

Supplementation of standard growth media (cation-adjusted Mueller-Hinton Broth [CAMHB]) with bicarbonate (**NaHCO**_**3**_) significantly increases β-lactam susceptibility of selected MRSA strains (“NaHCO_3_-responsive”). This “sensitization” phenomenon translated to enhanced β-lactam efficacy in a rabbit model of endocarditis. The present study evaluated NaHCO_3_-mediated β-lactam MRSA sensitization using an *ex vivo* pharmacodynamic model, featuring simulated endocardial vegetations (**SEVs**), to more closely mimic the host microenvironment. Four previously described MRSA strains were used: two each exhibiting *in vitro* “NaHCO_3_-responsive” or “NaHCO_3_-nonresponsive” phenotypes. Cefazolin (**CFZ**) and oxacillin (**OXA**) were evaluated in CAMHB±NaHCO_3_. Intra-SEV MRSA killing was determined over 72 hr exposure. In both NaHCO_3_-responsive strains, supplementation with 25 mM or 44 mM NaHCO_3_ significantly reduced β-lactam MICs to below the OXA susceptibility breakpoint (≤ 4 mg/L) resulting in bactericidal activity (≥ 3 log kill) in the model for both OXA and CFZ. In contrast, neither *in vitro*-defined NaHCO_3_-nonresponsive MRSA strains showed significant sensitization in the SEV model to either β-lactam. At both NaHCO_3_ concentrations, the fractional time-above-MIC was >50% for both CFZ and OXA in the NaHCO_3_-responsive MRSA. Also, in RPMI+10% LB media (proposed as a more host-mimicking microenvironment and containing 25 mM NaHCO_3_), both CFZ and OXA exhibited enhanced bactericidal activity against each NaHCO_3_-responsive strain in the SEV model. Neither CFZ nor OXA exposures selected for high-level β-lactam-resistant mutants within SEVs. Thus, in this *ex vivo* model of endocarditis, in the presence of NaHCO_3_ supplementation, both CFZ and OXA are highly active against MRSA strains that demonstrate similar enhanced susceptibility in NaHCO_3_-supplemented media *in vitro*.

## INTRODUCTION

*Staphylococcus aureus* is an important human pathogen that is associated with both community- and nosocomial-associated infections. It is a major cause of skin, soft-tissue, respiratory, and bone and joint infections, and is the leading cause of bacteremia and endocarditis in industrialized countries (1, 2). In many geographic locales, methicillin-resistant *Staphylococcus aureus* (MRSA) are responsible for the majority of *S. aureus* bacteremias (3), especially in community-associated scenarios (4). The Centers for Disease Control and Prevention (CDCP) has deemed MRSA (particularly community-acquired MRSA) as a critical “concerning threat” to human health (5).

Despite development of newer anti-MRSA antibiotics over the past decades (e.g., daptomycin; linezolid; oritavancin; dalbavancin and ceftaroline) (3, 6), these agents have been problematic for several reasons, including: cost; toxicities; lack of proven efficacy in bacteremia and endocarditis; and/or evolution of resistance during therapy, associated with clinical failures (7-9). Therefore, clinicians continue to rely on exploring novel approaches with older antimicrobials with proven efficacy, such as multi-drug combination treatments (10).

One of the limitations of antibiotic use is the predictive value of antimicrobial susceptibility testing and breakpoint determinations for selection. Although other diagnostics for infectious diseases have rapidly evolved over the past decade, antimicrobial susceptibility testing for MRSA has largely remained unchanged since the 1960s, with the use of bacterial growth medium such as Mueller-Hinton broth and agar (MHB; MHA). For MRSA susceptibility testing with oxacillin (OXA) to determine MRSA phenotype, the Clinical and Laboratory Standards Institute (CLSI) recommends growth of bacteria in 2% NaCl cation-adjusted (CA)-MHB (11, 12). Although this latter medium will amplify the capacity to isolate the OXA-resistant subpopulations within MRSA strains, it does not accurately reflect the host milieu; thus, the resulting minimum inhibitory concentrations (MICs) may not accurately represent antibiotic activity against MRSA within host-specific microenvironments (13).

Recent studies have focused on refining *in vitro* growth media in order to better simulate the host microenvironment *in vivo* in the context of more relevant and translatable antimicrobial susceptibility testing. Supplementation of standard media with bicarbonate (**NaHCO**_**3**_), a ubiquitous buffer in humans, has become the subject of several such studies (13, 14). These reports have demonstrated the ability of NaHCO_3_ to alter the susceptibility of selected MRSA strains *in vitro* to β-lactams. These investigations have focused on two conventional, prototype β-lactams not recommended for use against MRSA, OXA and cefazolin (**CFZ**) (14). These data enabled identification of two distinct MRSA phenotypes, NaHCO_3_-“responsive” and NaHCO_3_-“nonresponsive”. These *in vitro* phenotypes accurately predicted the ability of these same β-lactams to clear MRSA from multiple target tissues in a rabbit model of MRSA infective endocarditis (i.e., cardiac vegetations, kidneys and spleen) (14). The mechanism(s) by which NaHCO_3_ sensitizes “responsive” strains to such β-lactams appears to involve, at least in part, multiple genes which are critical in maintaining the MRSA phenotype, such as *mecA* and *sarA* (12); these perturbations may, in turn, lead to decreased production and/or maturation of PBP2a, yielding a “functional MSSA” phenotype (14).

The present study expands on our previous findings of bicarbonate sensitization of MRSA to β-lactams using a pharmacodynamic model featuring *ex vivo* simulated endocardial vegetations (**SEVs**). We hypothesized that ‘bicarbonate responsiveness’ in MRSA in this model would mirror similar findings in the rabbit endocarditis model, and that host-mimicking media within SEVs would help identify novel pharmacodynamic optimization strategies for prototypical β-lactams against such NaHCO_3_-responsive MRSA strains.

## RESULTS

### β-lactam susceptibilities in standard and alternative media supplemented with NaHCO_3_

CFZ and OXA MICs have been previously reported for these four study strains (14). In this investigation, we confirmed the strain-dependent NaHCO_3_ enhancement of the susceptibility of these isolates to the two study β-lactams. As noted in **Table 1**, strains 11-11 (USA300) and MW2 (USA400) displayed a significant reduction in β-lactam MICs in media supplemented with NaHCO_3_ (44 mM), while β-lactam susceptibility was not affected with NaHCO_3_ supplementation for the other two strains, COL (USA100) and BMC-1001 (USA500). In the two NaHCO_3_-responsive strains, supplementation with 44 mM NaHCO_3_ reduced the CFZ MIC to below the OXA susceptibility breakpoint (≤ 4 mg/L), with an 8-16-fold MIC reduction. Similarly, for these same strains, the OXA MICs were reduced 16-64-fold with 44 mM NaHCO_3_ supplementation.

**Table 1.** Cefazolin and oxacillin MICs in standard and alternative media types used in the pharmacodynamic model.

| Media | CFZ (mg/L) |  |  |  | OXA (mg/L) |  |  |  |
| --- | --- | --- | --- | --- | --- | --- | --- | --- |
|  | 11-11 | MW2 | COL | BMC-1001 | 11-11 | MW2 | COL | BMC-1001 |
| CAMHB-Tris<sup>a</sup> | 16 | 8 | 256 | 256 | 64 | 32 | 512 | 256 |
| CAMHB-Tris + 2%NaCl | 16 | 8 | 256 | 256 | 64 | 32 | 512 | 256 |
| CAMHB-Tris-NaCl + 25 mM NAHCO<sub>3</sub> | 4 | 8 | 256 | 256 | 4 | 32 | 512 | 256 |
| CAMHB-Tris-NaCl + 44 mM NAHCO<sub>3</sub> | 1 | 1 | 256 | 256 | 1 | 2 | 512 | 256 |
| RPMI 1640 + 10% LB | 1 | 2 | 32 | 8 | 1 | 1 | 64 | 4 |

<sup>a</sup>CAMHB-Tris was used only for susceptibility testing and not in the *ex vivo* SEV model since similar susceptibility in all strains occurred with addition of 2% NaCl.

It has recently been reported that an ‘antibiotic sensitization’ effect, especially for gram-negative bacteria and selected β-lactams, can occur in other host-mimicking media (13-15). The standard cell culture media, RPMI 1640, contains physiologic concentrations of NaHCO_3_ (∼25 mM). In this media, as opposed to CA-MHB, we noted substantially increased β-lactam susceptibility in all four strains regardless of genotypic background. However, the two NaHCO_3_-responsive strains were generally more responsive in this host-mimicking media, with resultant β-lactam MICs of ≤ 2 mg/L (**Table 1**), consistent with previous findings (14).

### β-lactam pharmacokinetics in the *ex vivo* SEV model

The pharmacokinetic profiles of antibiotics within the *ex vivo* SEV model is computer designed to precisely mimic actual patient exposures clinically. **Table 2** provides the predicted vs. observed pharmacokinetic profiles of CFZ and OXA in the SEV model’s central ‘fluid’ compartment (media). We simulated high-dose CFZ administration (2 g every 8 hrs) and OXA (2 g every 6 hrs), as recommended for invasive MSSA infections (16, 17). The concentrations achieved in the *ex vivo* model closely correlated to the targeted parameters for both antibiotics.

**Table 2.** Pharmacokinetics of CFZ and OXA in the central compartment (media) of the pharmacodynamic model. Data presented as mean ± standard error.

| Parameter | CFZ 2 g every 8 h |  | OXA 2 g every 6 h |  |
| --- | --- | --- | --- | --- |
|  | Predicted | Observed (n=12) | Predicted | Observed (n=12) |
| $C_{\max}$ (mg/L) | 256 | 249.4 ± 2.7 | 150 | 152.1 ± 1.4 |
| $C_{\min}$(mg/L) | 16 | 15 ± 1 | 2.3 | 2.2 ± 0.1 |
| $k_e$ (hr<sup>-1</sup>) | 0.385 | 0.354 ± 0.032 | 0.693 | 0.705 ± 0.005 |
| Half-life (h) | 1.8 | 2.0 ± 0.2 | 1 | 1.0 ± 0.0<sup>a</sup> |
| AUC<sub>0-24</sub> (mg/L*h) | 2442 | 2384 ± 31 | 1162 | 1217 ± 91 |

<sup>a</sup>standard error is <0.01

### Antimicrobial activity in the *ex vivo* SEV model

The antibiotic activities in the *ex vivo* SEV pharmacodynamic model are displayed in **Figures 1-3**; **Table 3** compares the ability of each β-lactam to reduce the MRSA counts within the SEVs over the 72 hrs course of treatment (expressed as the area under the bacterial curves, AUBC). As a point of reference, the less the AUBC, the more active the antibiotic regimen (18). The following trends emerged from these studies:

**Table 3.** Area Under the Bacterial Curve (AUBC) of CFZ and OXA treatment in standard and alternative media. AUBC is inversely related to antibiotic activity with lower AUBC indicating greater antibiotic effect. Data represent mean ± SD of duplicate replicates with two samples taken at each time point (n=4)

| Regimen | Media | AUBC |  |  |  |
| --- | --- | --- | --- | --- | --- |
|  |  | 11-11 | MW2 | COL | BMC-1001 |
| CFZ | CAMHB-Tris-NaCl (control) | 626.2 ± 6.6 | 658 ± 3.0 | 648.4 ± 2.8 | 633.9 ± 4.5 |
| | CAMHB-Tris-NaCl + 25 mM NaHCO<sub>3</sub> | 455.3 ± 4.8<sup>a</sup> | 489.2 ± 2.5<sup>a</sup> | 612.1 ± 2.5 | 592.7 ± 3.4 |
| | CAMHB-Tris-NaCl + 44 mM NaHCO<sub>3</sub> | 310.0 ± 10.5<sup>a,b</sup> | 390.8 ± 4.8<sup>a,b</sup> | 601.9 ± 4.5 | 566.7 ± 2.9<sup>a</sup> |
| | RPMI 1640 + 10% LB | 378.3 ± 3.5<sup>a,b</sup> | 387.3 ± 6.9<sup>a,b</sup> | 505.3 ± 8.1<sup>a</sup> | 538.8 ± 6.3<sup>a</sup> |
| OXA | CAMHB-Tris-NaCl | 610 ± 3.5 | 661.9 ± 9.5 | 621.1 ± 3.6 | 642.4 ± 5.1 |
| | CAMHB-Tris-NaCl + 25 mM NaHCO<sub>3</sub> | 480 ± 6.3<sup>a</sup> | 539.2 ± 23.3<sup>a</sup> | 596.7 ± 6.7 | 617.3 ± 2.6 |
| | CAMHB-Tris-NaCl + 44 mM NaHCO<sub>3</sub> | 392 ± 17.8<sup>a,b</sup> | 399.4 ± 15.0<sup>a,b</sup> | 568.9 ± 11.0<sup>a</sup> | 603.0 ± 5.38 |
| | RPMI 1640 + 10% LB | 415.1 ± 4.2<sup>a,b</sup> | 438.1 ± 9.3<sup>a,b</sup> | 649.7 ± 4.7 | 599.1 ± 9.4 |

<sup>a</sup>P<0.05 versus control
<sup>b</sup>P<0.05 versus 25 mM NAHCO<sub>3</sub>

**Figure 1.**
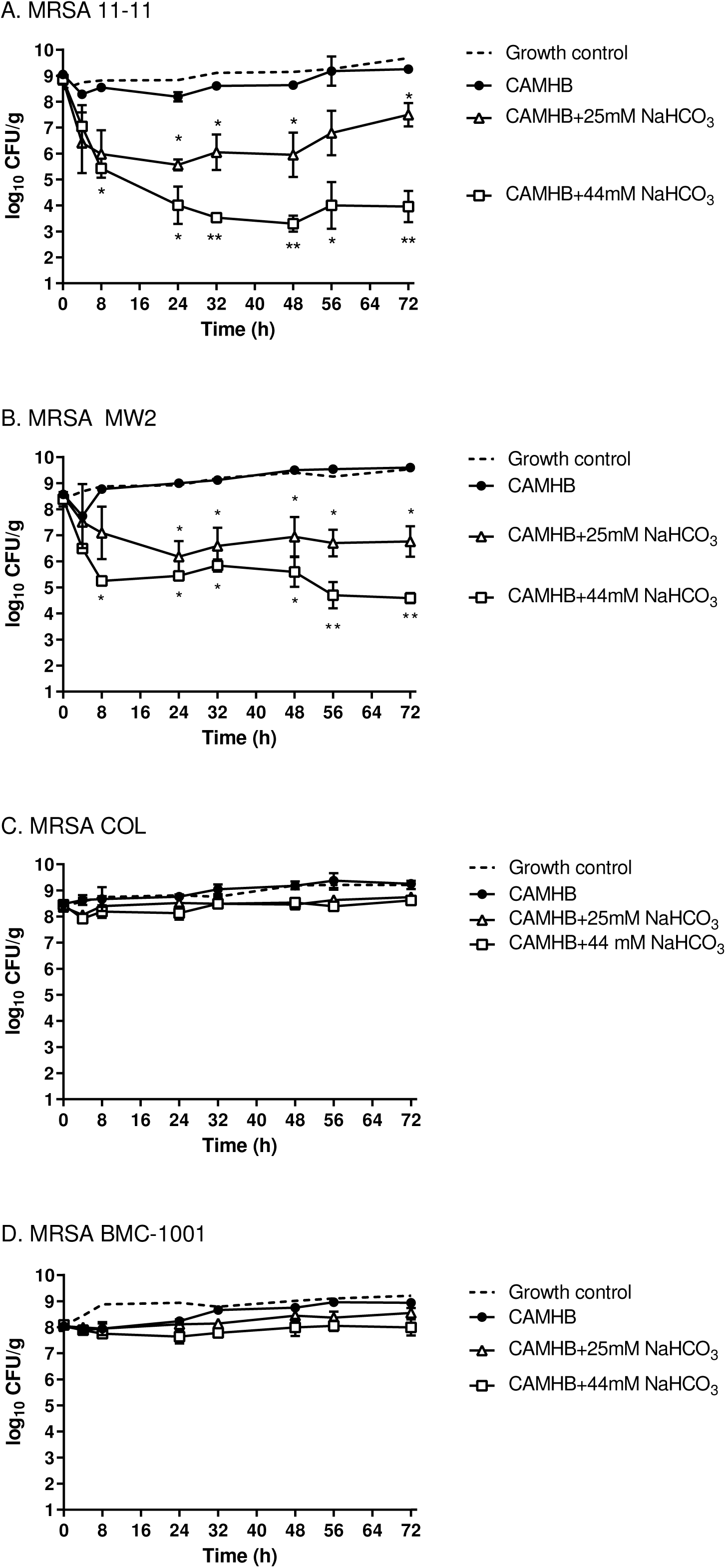
Kill curve activity of CFZ 2 g every 8 hrs simulations in the *ex vivo* SEV model in standard media, CAMHB-Tris-NaCl (represented as CAMHB), and NaHCO_3_-supplemented CAMHB against MRSA strains 11-11 (A), MW2 (B), COL (C), and BMC-1001 (D). Dashed line is untreated growth in CAMHB, solid lines indicate CFZ regimens. * = P < 0.05 vs CAMHB and ** = P < 0.05 vs CAMHB 25mM NaHCO_3_ media.

**Figure 2.**
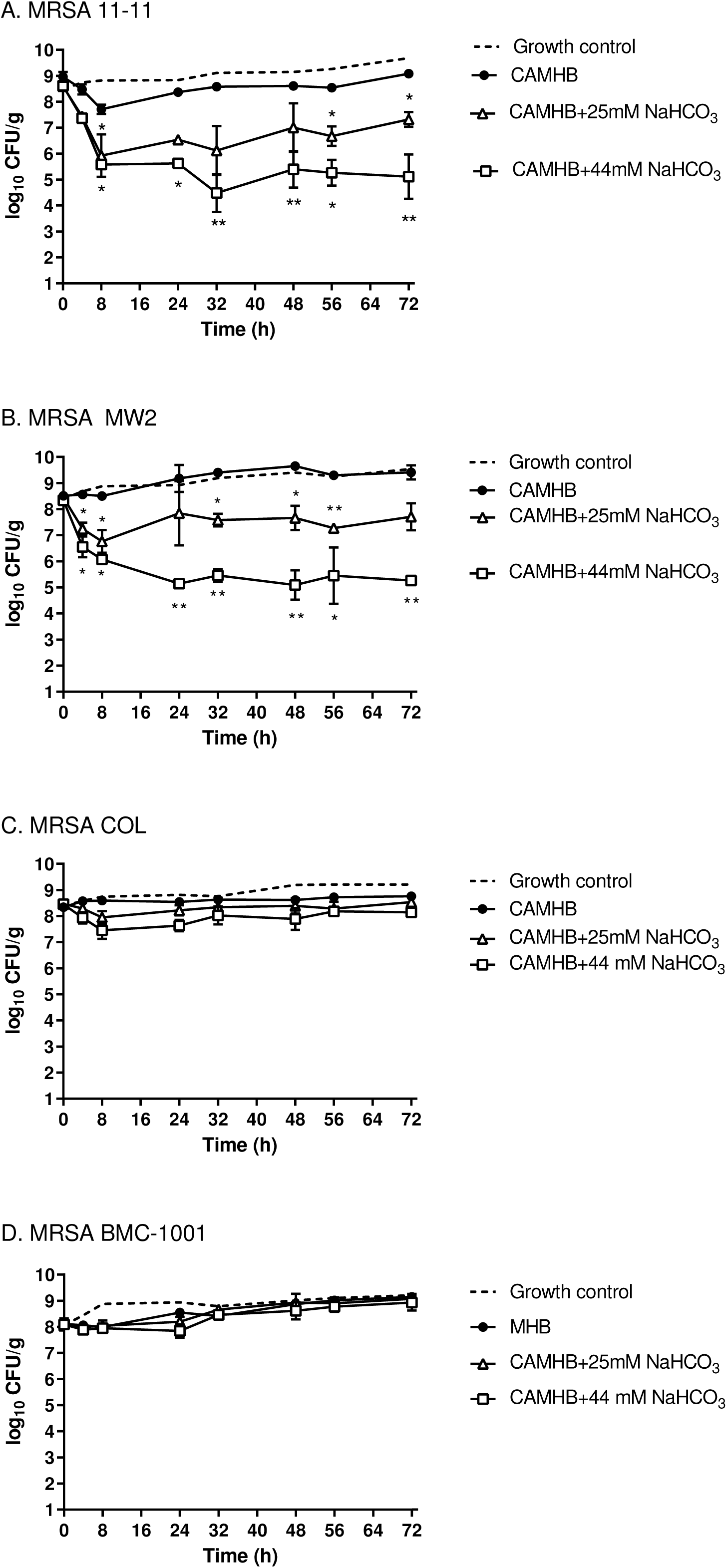
Kill curve activity of OXA 2 g every 6 hrs simulations in the *ex vivo* SEV model in standard media, CAMHB-Tris-NaCl (represented as CAMHB), and NaHCO_3_-supplemented CAMHB against MRSA strains 11-11 (A), MW2 (B), COL (C), and BMC-1001 (D). Dashed line is untreated growth in CAMHB, solid lines indicate OXA regimens. *= P < 0.05 vs CAMHB and **= P < 0.05 vs CAMHB 25mM NaHCO_3_ media.

**Figure 3.**
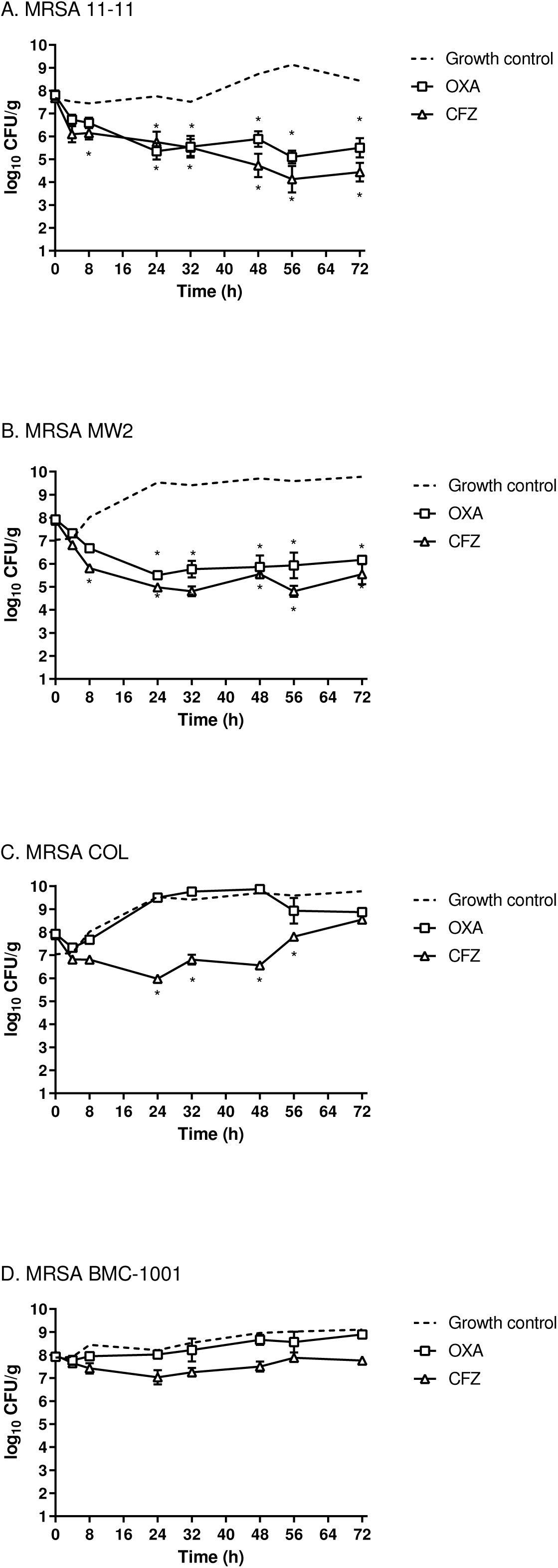
Kill curve activity of CFZ 2 g every 8 hrs and OXA 2 g every 6 hrs simulations in the *ex vivo* SEV model in host-mimicking RPMI+10%LB media against MRSA strains 11-11 (A), MW2 (B), COL (C), and BMC-1001 (D). *=P < 0.05 vs control.

- *β-lactams were inactive against all MRSA in the ex vivo SEV model with CAMHB in the central compartment without bicarbonate supplementation.* As expected, based on intrinsic MICs, neither CFZ nor OXA had effect against the four strains in the SEV model at human-equivalent, pharmacokinetic-based simulated dose-regimens. The activity curves among the strains were indistinguishable regardless of clonal background at all time points (**Figures 1;** as well as for the overall exposure based on similar AUBC values (**Table 3**). Based on the MICs determined in CAMHB, OXA achieved zero percent *ƒ*T>MIC for all strains, while CFZ achieved zero percent *ƒ*T>MIC for COL and BMC-1001 and 33% and 57% *ƒ*T>MIC for 11-11 and MW2, respectively.
- *Bicarbonate supplementation of CAMHB in the central compartment resulted in significant CFZ or OXA bactericidal activity in responsive (but not in non-responsive) MRSA strains in the ex vivo SEV model.* **Figure 1** displays the pharmacodynamic SEV kill curve of CFZ against all four strains with and without NaHCO_3_ supplementation. Based on MIC reductions noted in strains 11-11 and MW2 in this study, as well as in our previous *in vitro* work (14), we predicted that CFZ and OXA would each yield substantial intra-SEV antimicrobial activity against these responsive strains in the presence of NaHCO_3_ supplementation. Accordingly, in the *ex vivo* SEV model supplemented with NaHCO_3_, CFZ and OXA resulted in significantly greater anti-MRSA activity as compared to standard CAMHB medium in these responsive strains. After 8 hrs in the SEV model, β-lactam exposures in bicarbonate-supplemented media resulted in significantly greater activity as compared to standard CAMHB media (**Figure 1**, P<0.05 for timepoints 8-72 hrs). In comparing *ex vivo* kill curves between the different NaHCO_3_ concentrations, β-lactam exposure of the “responsive strains” in CAMHB medium supplemented with 44 mM NaHCO_3_ yielded a significantly greater bacterial count reduction for both CFZ and OXA vs 25 mM NaHCO_3_ supplementation at most time points from 24-72 hrs (**Figure 1 and Figure 2**, P<0.05). In contrast, there was no difference in killing of the nonresponsive strains, COL and BMC-1001, with NaHCO_3_ supplementation (vs either antibiotic-containing standard CAMHB medium or in untreated growth controls).

Overall, there was a notable NaHCO_3_-concentration response, with higher bacterial count reductions, faster time to bactericidal activity (time to 3 log_10_ CFU/g reduction), and lower AUBC when media was supplemented with 44 mM vs 25 mM NaHCO_3_ (**Figure 1; Table 3**. This correlated with greater susceptibility with the higher NaHCO_3_ concentration, resulting in 100% *ƒ*T>MIC for both the 11-11 and MW2 strains with 44 mM NaHCO_3_ vs 57% and 83% *ƒ*T>MIC, respectively, with 25 mM NaHCO_3_ for these strains. Since no change in MIC occurred in CAMHB + NaHCO_3_ in the nonresponsive strains, the *ƒ*T>MIC remained at zero percent, and reflects the lack of any β-lactam activity against those strains.

- *CFZ and OXA are each bactericidal ex vivo against NaHCO*_3_-*responsive (but not against non-responsive) MRSA in the host-mimicking medium, RPMI.* As noted above, CFZ and OXA MICs for all four strains were substantially reduced in RPMI. We next determined if these, *in vitro* outcomes were mirrored *ex vivo* in the SEV model. As displayed in **Figure 3**, the NaHCO_3_-responsive strains, 11-11 and MW2, were significantly killed with CFZ or OXA treatment in this medium; this is reflected in the enhanced pharmacodynamic attainment in this medium, resulting in 100% *ƒ*T>MIC for CFZ and 60% for OXA against both 11-11 and MW2 strains. In contrast, over the same 72 hrs β-lactam exposure period, the two nonresponsive strains (COL and BMC-1001) were minimally affected by either agent. It should be noted that there was initial activity with CFZ against COL in RPMI, with ∼2 log_10_ CFU/g killing at 24-48 hrs. However, this was not sustained after 48 hours, and the strain regrew to the initial inoculum. These data are reflected in the lower target attainment of 8.2%-56% *ƒ*T>MIC for CFZ and 0%-20% for *ƒ*T>MIC for OXA.
- *β-lactam treatment did not select for high-level CFZ or OXA resistance regardless of the NaHCO*_*3*_*-responsivity phenotype*. One additional advantage of this *ex vivo* model system is the ability to screen for emergence of drug-resistant mutants during human-simulated treatment strategies. In all our SEV simulations, we screened for evolution of high-level β-lactam-resistant mutants (≥ 4 x initial MIC) at time zero vs. every 24 hours thereafter. No such resistant variants were confirmed over the course of treatment with either CFZ or OXA.

## DISCUSSION

The β-lactam class of antibiotics remains the treatment of choice for a broad range of susceptible pathogens due to their potent and rapid mechanism(s) of action, as well as their relatively low rates of adverse side effects and toxicities. In addition, they have recently become well-characterized as able to augment the innate host immune system, via their synergistic interactions with host defense peptides against key gram-positive pathogens, including *S. aureus* (19-21).

The current study extends our previous work on the ability of NaHCO_3_ supplementation of standard MRSA *in vitro* media to identify a sub-set of MRSA strains that may, in fact, exhibit β-lactam susceptibility (14, 22, 23). The notion of employing alternative media for antibiotic susceptibility screening is not new; however, the biological basis and clinical utility of this approach has substantially increased in the last few years. For example, Kubicek-Sutherland *et al* mimicked the intracellular conditions of the macrophage phagosome in which *Salmonella* resides, using a mildly acidic and phosphate/magnesium-poor culture medium. Their study revealed that when *Salmonella* were grown in this media, antimicrobial susceptibility profiles emerging from this scenario were better predictors of treatment outcomes in murine bacteremia models than more traditional susceptibility testing strategies (23). Similar studies using *Acinetobacter* and staphylococci grown in Dulbecco Modified Eagles Medium (DMEM) or RPMI were better predictors of *in vivo* treatment outcomes, when compared to standard susceptibility testing modalities (13, 15). It should be emphasized that all four of our prototype MRSA isolates were rendered significantly more β-lactam-susceptible *in vitro* in RPMI media (4- to 64-fold MIC reductions). However, these salutary *in vitro* results in RPMI did not accurately predict subsequent microbiologic outcomes in the SEV model.

The enhanced antimicrobial activity seen above in such alternative media, as well as their outcome predictability *in vivo* may be, in part, reflective of: **i**) the enhanced activity of host defense peptides in combination with antibiotics *in vivo*; **ii**) bacterial adaptations *in vivo* that prevent excessive microbial growth; and/or **iii**) blunted expression of antimicrobial resistance mechanisms (14, 15, 20). However, it should be noted that some host-mimicking environments may actually be beneficial for bacterial survival. For example, under more acidic conditions, *E. coli* may become more resistant to β-lactams due to higher PBP1b production needed for organism survival in this harsh environment (24).

The treatment of MRSA infections poses several daunting challenges. Current treatment options for MRSA infections are relatively limited, and include antibiotics that are not only costly, but also substantially less effective, and often more toxic when compared to standard antibiotic treatments for MSSA (25-27). This is in large part due to the perceived inability to use traditional β-lactam antibiotics for MRSA infections (6). The *mecA* gene is primarily responsible for mediating *S. aureus* resistance to traditional β-lactams, encoding for penicillin-binding protein 2A (PBP2A), which has low affinity to most standard-of-care antistaphylococcal β-lactam antibiotics (6). Despite this, combination therapy featuring β-lactams plus anti-MRSA agents such as vancomycin and daptomycin have proven to be effective in selected patients with recalcitrant MRSA infections (19, 28-30). In addition, many MRSA strains which exhibit reduced susceptibility to vancomycin and daptomycin often demonstrate a paradoxical increase in susceptibility to β-lactams, a phenomenon known as the “see-saw effect”; this provides an additional scenario in which β-lactam agents may provide synergistic efficacy for MRSA killing (19).

Following our discovery of the NaHCO_3_ responsiveness of several prototype MRSA strains in terms of β-lactam hypersusceptibility *in vitro* (14), we recently screened a large collection of well-characterized MRSA strains (n = 58) for this same phenotype. We identified that ∼three-quarters of this cohort were CFZ-susceptible in NaHCO_3_-supplemented CA-MHB, while ∼one-third were OXA-susceptible in the same media (31). We have investigated the potential mechanisms by which NaHCO_3_ may cause such β-lactam hypersusceptibility *in vitro* and demonstrated that NaHCO_3_-supplemented media can down-regulate expression of the *mecA* gene, and subsequent PBP2a production, as well as by impacting several genes involved in maintenance of the MRSA phenotype, including *sarA* and *blaZ* (14).

The pharmacodynamic parameter best correlated with efficacy of β-lactam antibiotics is *ƒ*T>MIC (time-above-MIC of the free drug); this metric has been highly studied *in vivo* and *in vitro*, and is validated to predict improved clinical outcomes in β-lactam-treated patients (32-35). In most investigations, β-lactams demonstrate optimal activity with *ƒ*T>MIC in the range of 40-60% (32). Relevant to our study, we found that the lower MICs observed in NaHCO_3_-supplemented media for two of our prototype strains (11-11 and MW2) resulted in an “MSSA-like phenotype” improving the potential for target attainment for both CFZ and OXA. For the NaHCO_3_-responsive strains, CFZ achieved at least 83% *ƒ*T>MIC, resulting in a substantial and durable bactericidal activity in the *ex vivo* SEV model. Similarly, OXA, with a calculated ∼60% *ƒ*T>MIC in the two NaHCO_3_-responsive strains, also achieved good bactericidal activity in the *ex vivo* model.

The *ex vivo* simulated endocardial vegetation (SEV) model has been extensively used to pharmacokinetically and pharmacodynamically study the impacts of many antimicrobials against several clinical bacterial isolates in order to verify these interactions in a setting more akin to the host tissue microenvironment than *in vitro* assays (36, 37). In addition, this model has been shown to successfully recapitulate the microbiologic results generated in several *in vivo* animal models, including rabbit endocarditis (IE) (38). In the present investigation, we used this same *ex vivo* pharmacokinetic-pharmacodynamic model to more systematically evaluate the activity of CFZ and OXA against our four prototype MRSA isolates, in a host-mimicking microenvironment. Our current results in the SEV model provided an important “bridge” between *in vitro* and *in vivo* outcomes with these strains. *First*, the two MRSA that were significantly β-lactam/NaHCO_3_-responsive *in vitro* and *in vivo* (14), showed enhanced intra-SEV killing by CFZ and OXA in the presence of NaHCO_3_. In parallel, the two β-lactam/NaHCO_3_-nonresponsive strains (as defined *in vitro* and *in vivo*) were not substantially killed by CFZ or OXA *ex vivo. Second*, as seen *in vitro*, there appeared to be a NaHCO_3_ concentration-optimum for the β-lactam/NaHCO_3_ sensitizing phenotype for the two responsive strains (11-11 and MW2); thus, significant killing was seen *ex vivo* within SEVs both at 25 and 44 mM, but with a substantially greater bactericidal effect seen at the latter concentration. In contrast, the excellent *in vivo* clearance of both NaHCO_3_-responsive MRSA which occurs *in vivo* at NaHCO_3_ concentrations of 20-25 mM (14) suggests that additional host factors within cardiac vegetations or other target organs are in-play to synergistically kill MRSA [e.g., PMNs; platelets, host defense peptides; antibody, etc. (13-15)].

There remains a current debate over the most appropriate β-lactam to treat serious MSSA infections, with CFZ or OXA as the primary candidates (25, 39); most recent MSSA bacteremia studies appear to favor CFZ (16, 26, 27). Further, on balancing adverse events vs efficacy metrics, CFZ is generally regarded as a safer option, due to its decreased renal toxicity risk as compared to the antistaphylococcal penicillins (16, 27). However, there remains concern about CFZ efficacy in “high-inoculum” *S. aureus* infections (e.g., endocarditis) due to the potential for the undetected presence and induction of genes for Types A or C cephalosporinases that hydrolyze CFZ (39, 40). In our current study, CFZ appeared to be unaffected by the high inoculum within SEVs, at least for the NaHCO_3_-responsive strains, 11-11 and MW2, with excellent CFZ bactericidal activity obtained despite starting inocula ≥ 1×10^8^ CFU/g.

In conclusion, this study further validates the potential clinical translatability of the intriguing finding of β-lactam/NaHCO_3_ sensitization of selected MRSA strains *in vitro*. It will be important to extend our studies to even larger clinical MRSA cohorts (31). Ultimately, a pivotal clinical trial will be required to fully adjudicate the clinical utility of defining MRSA strains as β-lactam/NaHCO_3_-responsive in the clinical microbiology laboratory.

## ACKNOWLEDGEMENTS

This study was supported in part by a research grant from the National Institutes of Health (NIAID 1RO1AI146078-01) to A.S.B. and (NIAID 1RO1AI132627- to W.E.R.

## MATERIALS AND METHODS

### Bacterial strains, media and antibiotics

The strains used in this study were all clinical MRSA isolates, and represent diverse contemporary clonal genotypes (USA types) found worldwide in MRSA infections. These included MRSA 11/11 (USA300), MW2 (USA400), COL (USA100), and BMC-1001 (USA500). These strains are well-described in the literature and were recently utilized to define NaHCO_3_-responsivity *in vitro* (14). All strains were stored at –80°C in tryptic soy broth with 15% glycerol until thawed for use. Bacteria were cultured on MHA and incubated at 37°C in ambient air. Five liquid culture media were used for susceptibility testing comparisons including i) cation-adjusted Mueller Hinton Broth (CAMHB, Difco) with the addition of 100 mM Tris (hydroxymethyl-aminomethane; Fisher Scientific) to maintain pH at approximately 7.3 ± 0.1, ii) CAMHB-Tris supplemented with 2% NaCl; iii) CAMHB-Tris-NaCl supplemented with 25 mM NaHCO_3_; iv) CAMHB-Tris-NaCl supplemented with 44 mM NaHCO_3_; or iv) tissue culture medium, Roswell Park Memorial Institute (RPMI) 1640 (Fisher Scientific) supplemented with 10% Luria-Bertani (LB) broth; this latter medium contains ∼25 mM NaHCO_3_, as well as biotin, vitamin B_12_, and PABA, and in addition, the vitamins, inositol and choline. Media types ii-v were used for the *ex vivo* SEV model. The β-lactam antibiotics, OXA and CFZ, were purchased as analytical powders from Sigma-Aldrich (St. Louis, MO), and prepared fresh prior to each experiment according to the manufacturer’s protocols.

### Antibiotic susceptibility assays

CLSI guidelines (11) for microbroth dilution were used to determine antibiotic susceptibility (MICs) with modifications to the recommended media (12). Bacteria were grown overnight at 35°C on MHA, and resulting colonies were suspended in the different media listed above to the equivalent of 0.5 McFarland standard. This was further diluted 1:100 for a final inoculum of 5 x 10^5^ CFU/ml. Antibiotics were serially diluted 2-fold, and MICs were defined by standard metrics (11) and determined in triplicate on two separate days (n = 6 replicates).

### Pharmacodynamic model with e*x vivo* simulated endocardial vegetations (SEVs)

Each organism inoculum was prepared by spreading 0.5 McFarland standard on three MHA plates and incubating for 18-24 hrs. Bacteria were collected by scraping plates, and then adding to 5 ml CA-MHB in suspension. SEVs were prepared in 1.5 ml microcentrifuge tubes by mixing 50 µL of this organism suspension (final inoculum, 10^8^ CFU/g) with 0.5 ml of pooled human cryoprecipitate antihemophilic factor prepared from plasma (fibrinogen, von Willebrand factor, factor VIII, factor XIII and fibronectin) and 250,000 to 500,000 pooled platelets from human volunteer donors (UW Health Blood Bank, Madison, WI). These components were lightly vortexed, and a monofilament line added to each mixture along with 0.05 ml Bovine thrombin (5,000 units/ml). After gentle stirring, the resulting SEVs were removed from the tubes and then suspended within the central ‘fluid phase’ of this model. This preparation results in SEVs weighing ∼0.7 g and containing 3-3.5 g/dL of albumin and 6.8-7.4 g/dL of total protein (equating to human physiologic levels). The protein binding of the study drugs has been found to be 84% for CFZ and 92% for OXA, which was used to calculate free AUC (*ƒ*AUC) and percent time of free drug above the MIC (%*ƒ*T>MIC) (41, 42).

The central (‘fluid’) compartment model for the SEVs consists of a 150 ml flask glass with sealed screw cap. The flask was prefilled with media and magnetic stir bar, and SEVs were added for 30 minutes prior to antibiotic dosing to allow for climate acclimation. The model was maintained at 35-37°C ambient air and fresh media instilled via a continuous syringe pump system (New Era Pump Systems, Inc.) to provide a human pharmacokinetic simulation of the antibiotics. All model experiments were performed in duplicate flasks to ensure reproducibility with two SEVs collected for each time point (n=4 SEVs per time point).

- *Simulated antimicrobial regimens*.All regimens were derived from human pharmacokinetic data and standard dosing regimens for humans with MSSA infections previously published: OXA = 2 grams infused every 6 hrs (43) or CFZ = 2 grams infused every 8 hrs (43, 44). Antibiotics were administered as boluses over 1 minute into a luer-lock port of the flask at the scheduled administration time, over 72 hrs dosing duration. The predicted pharmacokinetics of each regimen are provided in **Table 2**.

### Pharmacokinetic analysis and exposure determination

Pharmacokinetic samples were obtained in duplicate through the injection port of each model from 0-72 hrs for verification of target antibiotic concentration attainment. All samples were stored at -80°C until ready for analysis. Concentrations of OXA were determined by bioassay using *Kocuria rhizophila* ATCC 9341 on MHA as previously described (44). Concentrations of CFZ were determined by bioassay using test organism *Bacillus subtilis* ATCC 6633 on MHA (45). The half-lives, area-under-the-curve 0-24 hrs (AUC), maximum and minimum concentrations (C_max_, C_min_) of each antibiotic were determined by the trapezoidal method utilizing Prism (GraphPad Software, Inc.). Observed *ƒ*AUC/MIC and %*ƒ*T>MIC were determined in Prism and reported for OXA and CFZ.

### Assessment for emergence of variants with high-level β-lactam resistance in the *ex vivo* model

Samples of 100 µl from each time-point were parallel-plated onto MHA plates containing either no antibiotic or 4-fold the respective β-lactam initial MICs to assess for the emergence of high-level resistant mutants. Plates were then examined for growth after 48 hrs incubation at 35°C. Specific CFZ or OXA MICs were then performed on selected colonies exhibiting growth on the respective antibiotic-containing agar plates to quantify the actual changes in MIC over the 72 hrs β-lactam exposure period.

### Statistical analysis

Bacterial counts, expressed as log_10_ CFU/g, in SEVs at each time point were determined for each antibiotic treatment and growth condition for each strain. AUBC, defined as area under the bacterial growth curves (AUBC) over the 72 h experiments, was also calculated. A two-way analysis of variance was used with Tukey’s Post-Hoc test to compare bacterial counts and AUBC with a *P* value of ≤ 0.05 for significance. All statistical comparisons were analyzed using Prism 8 (GraphPad Software, San Diego, CA).

